# KhufuEnv, an auxiliary toolkit for building computational pipelines for plant and animal breeding

**DOI:** 10.1101/2025.03.28.645917

**Authors:** Hallie C. Wright, Catherine E. M. Davis, Josh Clevenger, Walid Korani

## Abstract

In the era of short and long read sequencing, vast amounts of DNA sequencing data are being generated. While a variety of tools exist for analyzing and manipulating genomics data, many have a finite number of tools, and thus, require users to depend on multiple sources for conducting analyses and processing data. An integrative environment of tools which is accessible to users of different computational backgrounds would facilitate more efficient data processing and level the playing field for researchers whose research depends on analyzing genomic data. We developed the KhufuEnv, an open-source, flexible, auxiliary environment for manipulating and analyzing genomic data, among other datasets, in the Unix environment. The KhufuEnv provides a buildable platform for constructing custom pipelines for several genomic analyses across different species. As a proof of concept, we demonstrate rapid *de novo* identification of previously characterized quantitative trait loci (QTL) and calculate the proportion of the genome containing runs of homozygosity (ROH) with in-house tools on published datasets. Additionally, we introduce our custom HapMap and PanMap file formats. The KhufuEnv can be exploited for a variety of applications and implemented for quick analysis, supporting users with minimal computational experience.

## Introduction

The emergence of DNA sequencing technology has brought forth a plethora of data. The National Center for Biotechnology Information (NCBI) sequence read archive (SRA) contains over 36 petabytes (PB) of sequencing data to-date (NCBI 2020). In fact, genomics data is predicted to be the most demanding domain of data relative to other big data generators like astronomy or YouTube (Stephens et al. 2015). Such data has revolutionized the capabilities of breeding programs and supported genomic selection (Meuwissen et al. 2001) in animals (Wiggans et al. 2017) and plants (Crossa et al. 2010). The wealth of data being generated, however, demands the ability to analyze it in a high-throughput and efficient manner (Bayat 2002). Thus, user-friendly tools which can support researchers with various computational skillsets are imperative for extracting relevant information from the masses of data being generated.

Several suites of tools have been developed for manipulating genomics data, including SeqKit (Shen et al. 2016; Shen et al. 2024), Bioawk (Li 2017), Seqtk (Li 2013), SeqFu (Telatin et al. 2021), VCFTools (Danecek et al. 2011), BBTools (Bushnell et al. 2017), and BEDTools (Quinlan and Hall 2010). Some, such as VCFTools, Seqtk, and SeqFu, operate on limited data formats (VCF and FASTA/FASTQ files, respectively). Others, like Bioawk, focus primarily on parsing column and header information from genomic data files. Although these programs have immense value in their own respects, many of them require users to exploit additional software to extract biological meaning. We present KhufuEnv, aimed to design a versatile and expansive environment for genomics data manipulation and trait mapping based on three principles: it requires simple processing, runs on a single thread, and can be completed quickly. To our knowledge, this is the largest collection of computational genomics tools within a single environment to-date.

Visually similar to the shape of a quantitative trait locus (QTL) peak (Fig. 1A), the Khufu platform (https://www.hudsonalpha.org/khufudata/), developed by Korani et al. (2021), was named after the Great Pyramid of Khufu (Bartlett 2014). In their 2021 study, Korani et al. demonstrated Khufu’s utility for conducting QTL-mapping with low-coverage sequencing of whole populations to *de novo* identify QTL for blanchability, which were validated in an independent population. This same platform also confirmed *de novo* QTL identified in previously described datasets (Chu et al. 2019; Clevenger et al. 2018).

**Figure 1:**
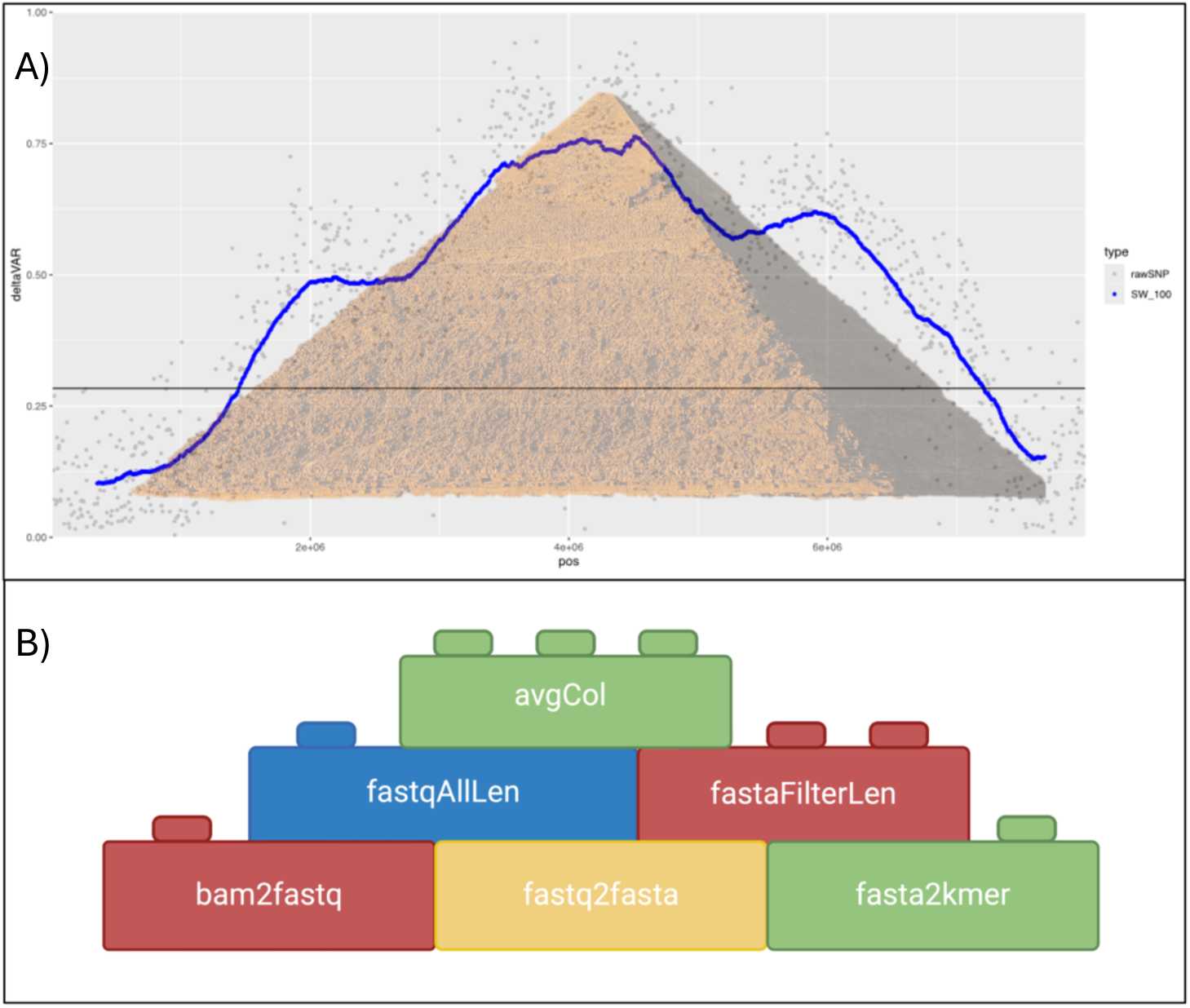
A) Inspired by the visualization of QTL mapping peaks and subsequently named after the pyramid of Khufu, the KhufuEnv is an extension of the Khufu platform described by Korani et al. 2021. B) The KhufuEnv is an auxiliary environment containing modular tools that facilitate building customized pipelines.

Although Khufu was originally conceived for mapping traits in complex polyploids like peanut (*Arachis hypogaea*), the KhufuEnv, an extension of the Khufu platform, contains versatile tools which can be exploited across a variety of plant, animal, human, and microbial systems.

The KhufuEnv is an auxiliary environment containing tools which support analyses of high throughput sequencing data. Tools in the KhufuEnv can be implemented modularly to construct customized pipelines (Fig. 1B). Here, we summarize the core sections of the KhufuEnv, introduce our custom HapMap and PanMap formats, and detail four case studies demonstrating the efficiency and utility of the tools in this environment.

## Overview

The KhufuEnv can be downloaded from https://github.com/w-korani/KhufuEnv, and the only dependencies required are R (R CoreTeam 2021) and gawk. The KhufuEnv contains six sections with 132 tools for manipulating genomics files and building custom pipelines. The sections include: KhufuEnvHapMap, KhufuEnvPanMap, KhufuEnvStats, KhufuEnvFASTA, KhufuEnvFASTQ, and KhufuEnvDataset. The following overview is not exhaustive of all tools. After one loads the environment, the command “KhufuEnvHelp” can be executed to list all tools. More detail regarding individual tools can be retrieved by running the command “KhufuEnvHelp [tool name].” For instance, to learn more about how to use the tool vcf2HapMap, one could run ***KhufuEnvHelp vcf2HapMap*** and obtain information regarding the tool’s description, parameters, and example commands associated with practice files located in the KhufuEnv (Supplementary Fig. 1).

### KhufuEnvHapMap

Variant calls are housed in a variety of file formats. Two popular formats are VCF (Danecek et al. 2011) and HapMap (The International HapMap Consortium 2003) files. In the KhufuEnv, we have created a modified version of the “standard” HapMap (Supplementary Fig. 2) which contains the most essential information regarding variant calls (Supplementary Fig. 3). The Khufu HapMap includes columns that correspond to chromosome, position, and sample names for genotyping calls at their respective sites. The Khufu HapMap has dropped columns for reference SNP cluster identifier (rs#), alleles, strand, reference assembly version (assembly#), genotyping center (center), protocol identifier (protLSID), genotyping assay identifier (assayLSID), panel of individuals genotyped (panelLSID), and quality control for entries (QCcode). Such information were perhaps more relevant to human studies conducted within the International HapMap Project (Thorisson et al. 2005) yet are trivial to the actual calling of variants. Allele calls with the value “NA” are converted to “-,” and homozygous allele calls are condensed to a single allele; i.e., “AA” is converted to “A”. Heterozygous allele calls still maintain information for two alleles. Henceforth in this paper, HapMap will refer to the Khufu HapMap format.

The KhufuEnvHapMap section contains several tools for processing HapMap files.

Genotyping files can be easily converted between text, VCF, standard HapMap, and HapMap. Note that values for filter, quality, and depth of coverage will not be present when converting HapMap to VCF since this information is not included in HapMap format. Often in genomic analyses, calls must be filtered to achieve a set of high confidence variants by removing potentially false variants. Within the KhufuEnv, HapMap variants can be filtered for those which are diallelic, single nucleotide polymorphisms (SNPs), or structural variants (SVs) and meet specified missing data and minor allele frequency (MAF) criteria. Similarly, the KhufuEnv allows for filtering out samples which have a percentage of missing variant calls.

### KhufuEnvPanMap

Visualizing long and short variants side-by-side can be a challenge using traditional file formats. For instance, VCFs may display large strings of DNA to represent a long SV, which is challenging for visualizing alleles on a command line or external (e.g. Excel) interface. We resolve such issues with the development of our custom PanMap format (Fig. 2). The PanMap format is similar to the HapMap format in that we retain information regarding chromosome, position, and calls for samples. However, the PanMap format also includes columns that list the length of variants called across parents and the allele call for each parent (designated numerically). Sample allele calls are called numerically with respect to parentals codes, and heterozygous sample calls include a comma to separate the alleles. Like the KhufuEnvHapMap section, there are several tools for processing PanMap files. PanMaps can be converted to and from HapMap, VCF, and heatmaps, and PanMap can be converted to text. PanMaps can also be filtered for samples with missing data, variants with missing data, and MAF.

**Figure 2:**
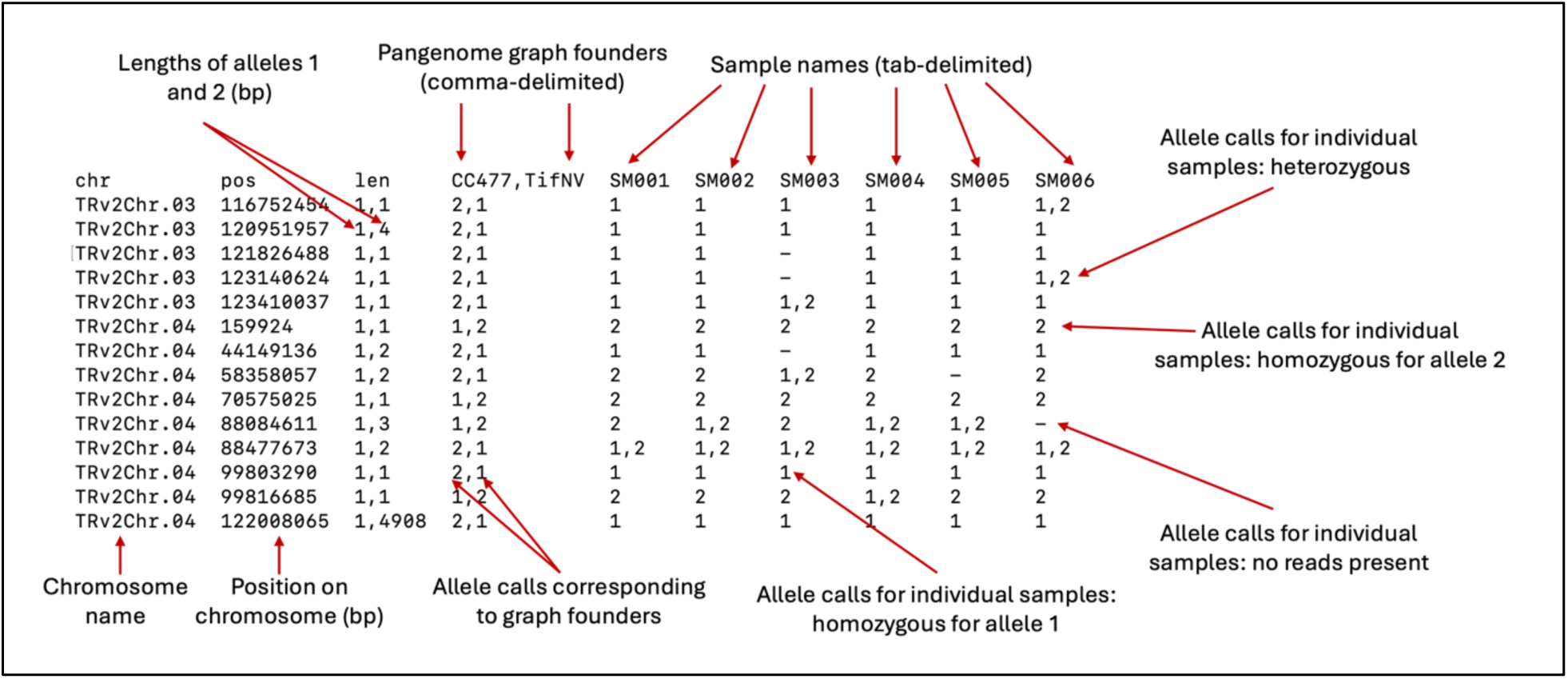
The Khufu PanMap file format concisely displays variants against a pangenome graph, outlining both short and long variants side-by-side. Arrows designate descriptions for different parts of the file.

### KhufuEnvStats

The KhufuEnvGeneralStats section covers tools that allow for easy extraction of statistics including average, median, standard deviation, sum, and minimum and maximum values for factors, columns, or rows in a dataset.

### KhufuEnvFASTA

This section contains tools to facilitate converting FASTAs to text files with fasta2txt and vice-versa with txt2fasta. Files can be split into individual chromosomes with fastaSplit, and sequences within a single file that have the same header can be combined in the same file with fastaCombine. Sequence length for each chromosome can be extracted with fastaSeqLen, and the entire sequence length can be extracted with fastaLen. FASTA files can be artificially k-merized with fasta2kmer, specifying window width and step size.

### KhufuEnvFASTQ

Like some of the tools in the KhufuEnvFASTA section, the KhufuEnvFASTQ section allows for manipulating FASTQ files which can aid in preprocessing DNA sequence data prior to analysis. FASTQs can readily be converted to FASTA and vice versa with the fastq2fasta and fasta2fastq tools. The entire length of nucleotides can be extracted with fastqAllLen, which is useful for calculating depth of coverage without prior knowledge of the sequencing depth for FASTQ files. The fastqLen tool can be used to print the length of each read individually from a FASTQ file, and the fastqAverage tool can be used to print the average length of reads. FASTQs can be filtered by minimum read length with the fastqFilterLen tool, subsampled for a desired total nucleotide length with fastqSubSampling, and artificially k-merized with fastq2kmer. The phred33 and phred64 tools print quality score information for the user’s reference.

### KhufuEnvDataset

The KhufuEnv is not exclusive for facilitating analysis of genomics data but can be extended to several other data types in the KhufuEnvDatasetProcessing section. Similar tools to those presented in the HapMap and PanMap sections, merge, subtract, and combineSimilarColumnDS can be applied to other datasets here. This section also includes tools which may be relevant to preparing datasets for statistical analyses such as long2wideDS and wide2longDS and extracting metadata for data sets tools like dimDS, maxLenDS, and minLenDS.

## Usage Examples and Common Scenarios

To demonstrate the utility of KhufuEnv for building custom analysis pipelines, we have provided four case studies related to plant and animal breeding using publicly available data. All case studies were run in the HudsonAlpha Institute for Biotechnology high performance computer (HPC) environment on a Rocky Linux operating system. Each task ran on less than 2.7 GB of memory.

### Case study 1) Confirming a candidate locus for cold tolerance in peanut (*Arachis hypogaea* L.)

Peanut (*Arachis hypogaea* L.) is a globally important oilseed and food crop. While it thrives in tropical and subtropical regions, it is cultivated in over 100 countries under a variety of agroecological environments (Variath and Janila 2017). The increasing demand for peanut (Das and Brown 2024) requires the expansion of production to high-latitude areas (Zhang et al. 2019). However, cold temperatures negatively impact peanut development (summarized by Zhang et al. 2019) and can be particularly detrimental to germination (Hurdle et al. 2020). To aid marker-assisted breeding for chilling tolerance in peanut, Zhang et al. (2023) identified a QTL for cold tolerance during germination in a recombinant inbred line (RIL) population from a bi-parental cross of a cold-susceptible (DF12) and cold-tolerant (Huayu 44) parent. The cold-susceptible and cold-tolerant parents were sequenced to a depth of coverage of 22.27x and 18.7x, respectively, and 200 random RILs were sequenced to an average depth of coverage of 5.76x (NCBI SRA accession number PRJNA931845). We exploited the KhufuEnv to demonstrate the rapid *de novo* identification of this QTL using this the dataset by Zhang et al. (2023) (Fig. 3).

**Figure 3:**
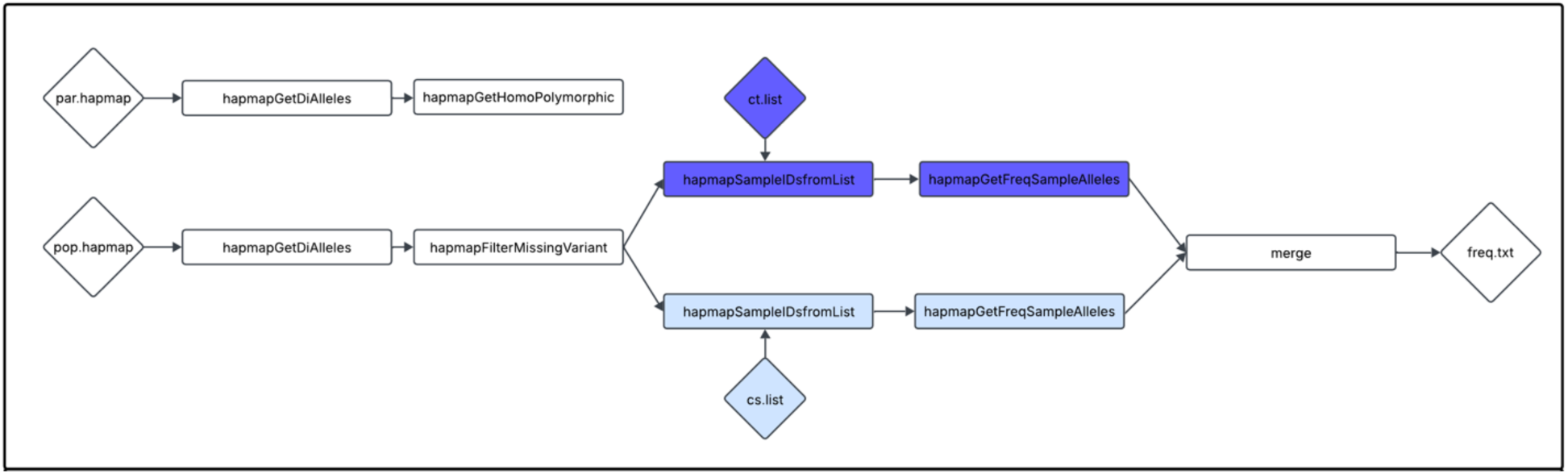
The custom KhufuEnv pipeline used for identifying a cold tolerance QTL in peanut.

A parental HapMap for the cold-susceptible (DF12) and cold-tolerant (Huayu 44) parents was constructed by using the Khufu pipeline described by Korani et al. (2021) (https://www.hudsonalpha.org/khufudata/) which exploits BWA (Li and Durbin 2009) to align reads from both samples to the TifRunner version 2 peanut assembly (https://data.legumeinfo.org/Arachis/hypogaea/genomes/Tifrunner.gnm2.J5K5/) and generates variant calls with a minimum read depth per variant of five. The hapmapSNP2SV tool was used to reformat the HapMap to distinguish SVs from SNPs with commas. The variants in the parental HapMap were then filtered to retain those which were diallelic with hapmapGetDiAlleles and homopolymorphic with hapmapGetHomoPolymorphic. A text file containing relative germination rates from Zhang et al. 2023 for each line across five independent experiments was used to extract 20 samples with the highest germination rates as cold-tolerant (ct) and the 20 samples with the lowest germination rates as cold-susceptible (cs). The sample names were used to generate ct and cs list files.

A population HapMap was generated by using the Khufu pipeline as described above with a minimum read depth per variant of three. The population HapMap was filtered to include only diallelic variants with hapmapGetDiAlleles and remove variants with > 50% missing data with hapmapFilterMissingVariant. Using two text files which contain the names of cs and ct samples, the population HapMap was split into separate cs and ct HapMaps with hapmapSampleIDsfromList. The hapmapGetFreqSampleAlleles tool was used to obtain allele frequencies, and each of the frequencies for the cs and ct samples were combined into a single text file using the merge tool. This allele frequency text file was processed to calculate Δ (SNP-index), i.e., the difference in allele frequencies across the two populations, and extract candidate regions which exhibited the highest Δ (SNP-index) across all environments. The highest Δ (SNP-index) is located near 155.2 Mb on chromosome 19 (B09) (Fig. 4), located within the same region that Zhang et al. (2023) fine-mapped the QTL to.

**Figure 4:**
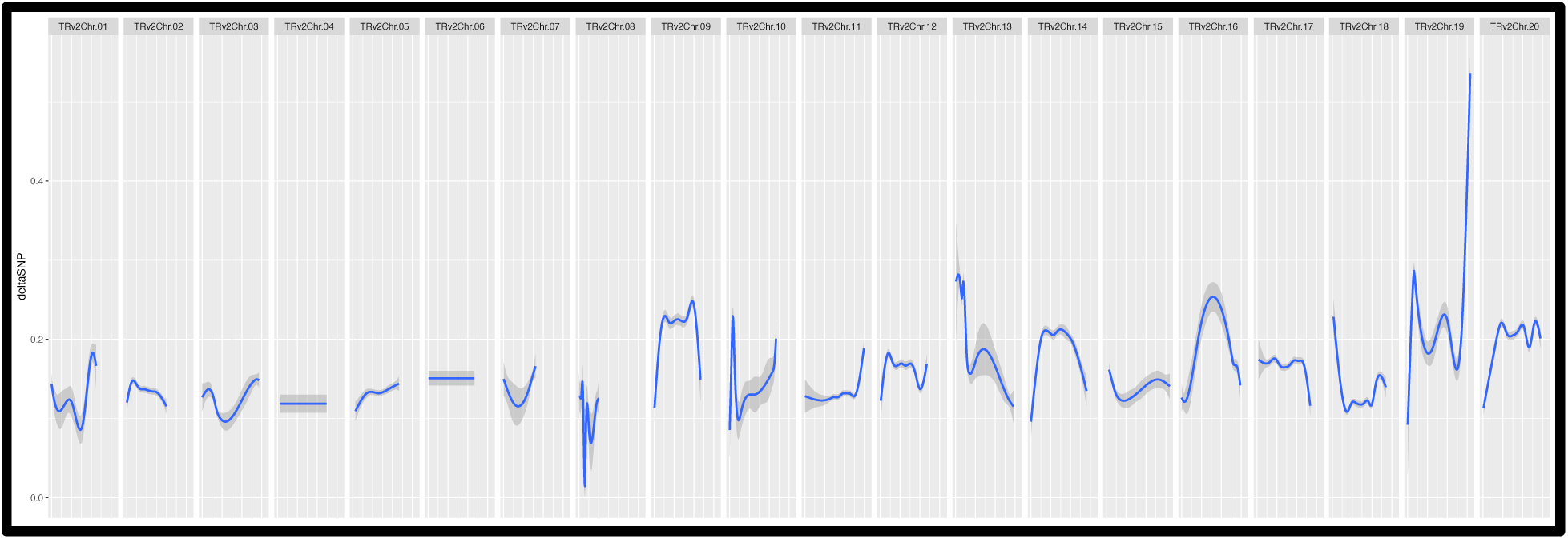
Tools in the KhufuEnv enable the calculation for the Δ (SNP-index) for 20 ct and 20 cs samples across the peanut genome, which demonstrate a strong association near 155.2 Mb on Chr19 for cold tolerance at germination.

### Case study 2) Confirming a sex-determination region (SDR) previously identified in *Amborella trichopoda*

The transition from monoecy, the presence of male and female organs on the same plant, to dioecy, the presence of male and female organs on separate plants, has independently evolved hundreds of times in flowering plants (Renner 2014). Dioecy is estimated to occur in 5-10% of flowering plants and is the result of genetically controlled sex determination systems (Renner 2014). SDRs are genomic regions which rarely recombine and contain sex-determining genes (Charlesworth 2002). Characterizing sex chromosomes and SDRs in plants is relevant for understanding genome evolution and has implications for plant breeding, particularly for fruiting crops (Charlesworth and Harkess 2024).

As a sister to all extant flowering plants (Soltis et al. 2011) and a dioecious plant (Anger et al. 2017), *Amborella trichopoda* is an ideal system to study sex chromosome evolution in flowering plants (Adams 2013; Soltis et al. 2011). Recent studies have shed light on its ZW sex determination system (Soltis et al. 2011), but the genes involved in dioecy in *Amborella* have previously remained unknown. Carey et al. (2024), however, recently characterized the SDR and identified candidate genes for sex determination in *Amborella*. We use the whole-genome sequencing (WGs) data for 104 *Amborella* samples (NCBI SRA accession number PRJNA1161132) and the Santa Cruz 75 version 2 assembly published by Carey et al. (2024) to demonstrate the KhufuEnv capabilities in identifying trait-associated regions in allosomes (Fig. 5).

**Figure 5:**
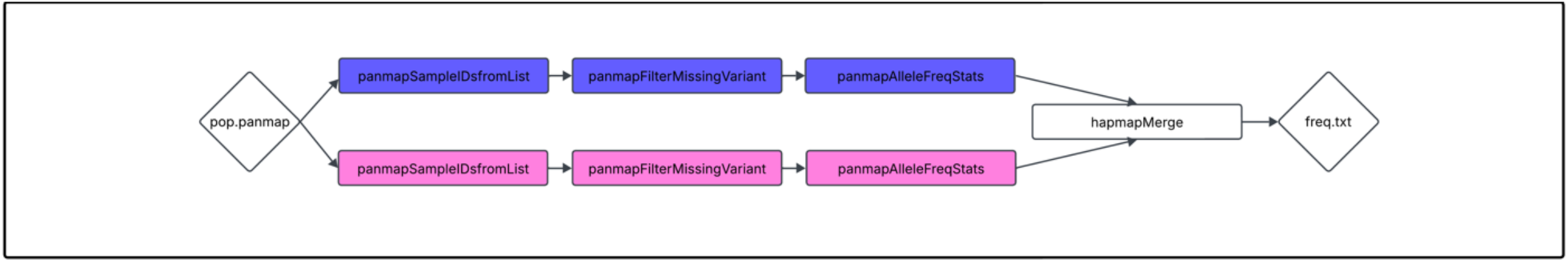
The custom KhufuEnv pipeline used for confirming previously identified SDR in Amborella.

Briefly, the KhufuPAN pipeline (Lee et al. 2025) (https://github.com/w-korani/KhufuPAN) was implemented to generate a pangenome for HAP1 and HAP2 of the Santa Cruz 75 assembly in the form of a graphical fragment assembly (GFA) file with Minigraph-Cactus (Hickey et al. 2024). After retaining high quality variants for the pangenome parents, Giraffe (Sirén et al. 2021) was used for mapping reads from the *Amborella* samples to the pangenome. A variant file in the form of a PanMap was generated using a minimum depth filter of two reads per variant. Initially, two list files were generated corresponding to male and female individuals described by Carey et al. (2024) and panmapSampleIDsfromList was used to generate separate male and female PanMaps. Each PanMap was filtered to remove variants with data missing from > 75% samples with panmapFilterMissingVariant.

Allele frequencies were obtained for each PanMap with panmapAlleleFreqStats. The column information for the chromosome, position, variant length, and parental calls from the PanMap was added to the allele frequencies to generate female and male allele frequency statistics files. The male and female allele frequency statistics files were then merged using hapmapMerge. The merged file was manually filtered to remove redundant parental call and length columns and columns containing allele count and sum.

Differences in allele frequencies which differ by > 0.75 were mined for putative candidate regions for SDR (Supplementary File 2). A single region between 44,825,527-47,193,113 bp on chromosome 9 contained allele frequencies which differed by 1.0 (Fig. 6A). This region was 245,680 bp downstream to the *MTN1-2* gene Carey et al. (2024) identified as a candidate sex-determining gene (Fig. 6B).

**Figure 6:**
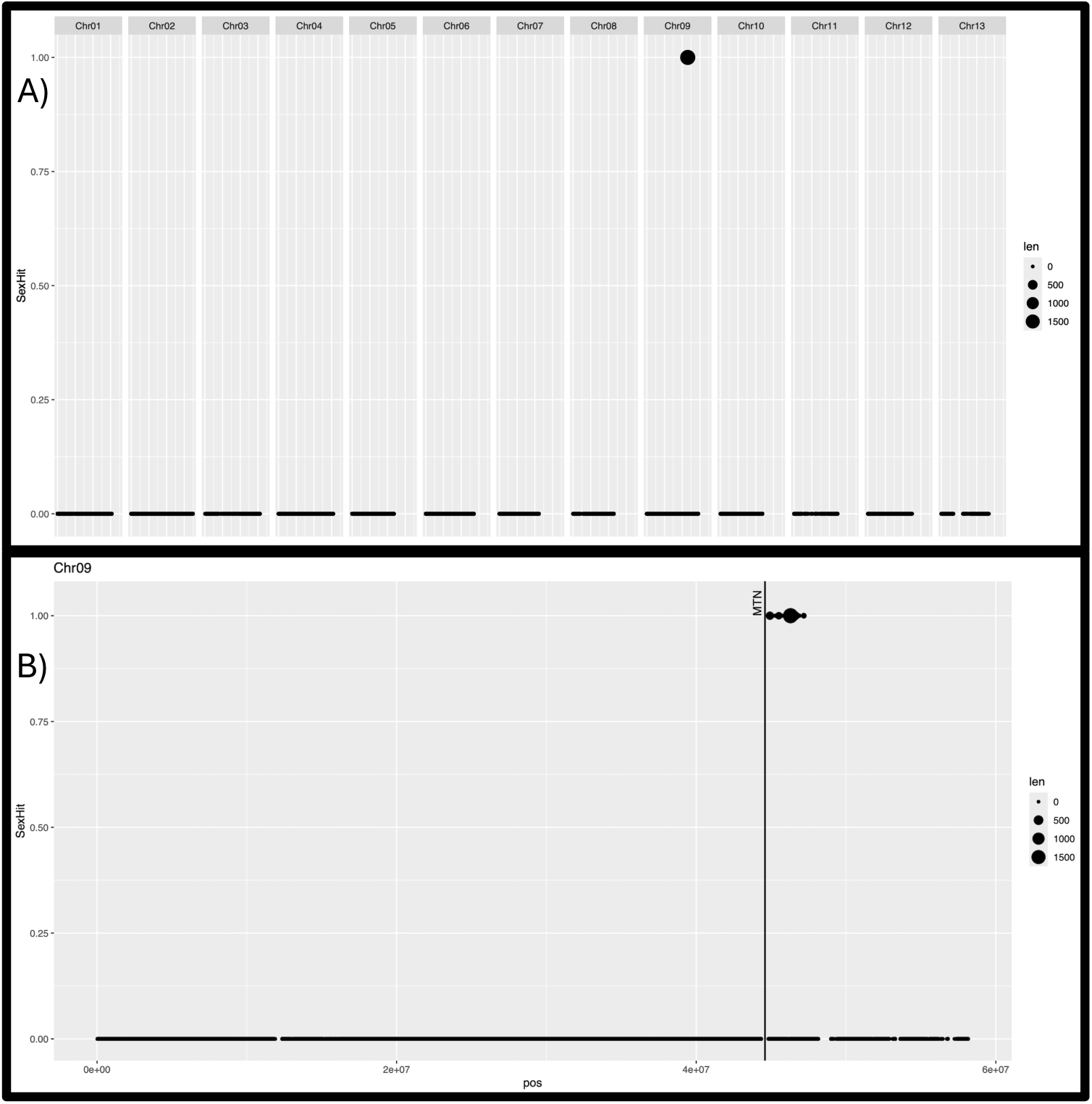
A) A single SDR on chromosome 9 contained allele frequencies which differed by > 0.75 between male and female WGS samples. B) The SDR identified with a custom Khufu pipeline was 246 Mb downstream to MTN1-2, a candidate sex-determination gene identified by Carey et al. 2024. “Len” represents the maximum length of an allele variant at a respective location.

### Case study 3) Estimating runs of homozygosity (ROH) in canine

Runs of homozygosity (ROH) (Broman and Weber 1999) are long stretches of homozygous regions resulting from inheritance of two copies of an ancestral haplotype. ROH can be used in animal breeding as measure of inbreeding within individuals or populations (Ferenčaković et al. 2013), and consequently, can be an indicator of selection (Mastrangelo et al. 2018; Zhang et al. 2015). In canines, ROH varies greatly across and within breeds in part due to inbreeding associated with selection (Anger et al. 2017; Sams and Boyko 2019). Greater ROH may be particularly alarming given that stretches of homozygosity often harbor deleterious alleles associated with disease (Subramanian and Kumar 2024) and may be an artifact of recent canine inbreeding (Anger et al. 2017). Meadows et al. (2023) conducted short read sequencing on nearly 2,000 candid samples at 20x depth of coverage to assess how ROH varies across over 300 dog breeds, village dogs, wolves, and coyotes. While Meadows et al. (2023) and others (Gorssen et al. 2021; Kim et al. 2024; Meyermans et al. 2020; Mulim et al. 2022; Peripolli et al. 2017) calculate ROH using PLINK (Purcell et al. 2007), we present a method for quickly calculating ROH and *F*_*ROH*_ in the KhufuEnv (Fig. 7).

**Figure 7:**
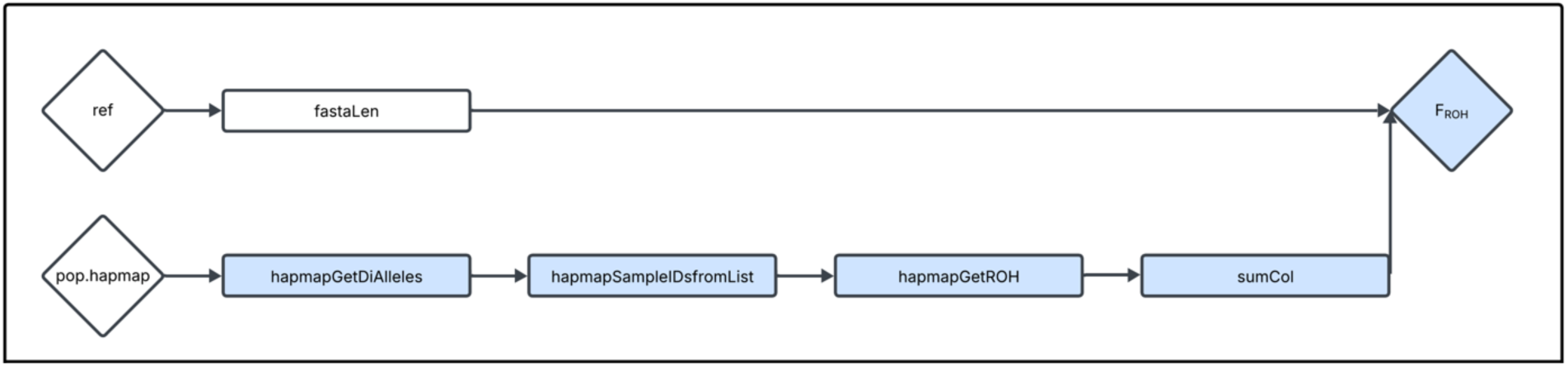
The custom KhufuEnv pipeline used for calculating *F*_*ROH*_ in canine.

To calculate *F*_*ROH*_, we first used the fastaLen tool to determine the total sequence length of the German Shepherd reference genome (Wang et al. 2021) (UU_Cfam_GSD_1.0, GCF_011100685.1). The variant calls published by Meadows et al. (2023) in the form of a 6 VCF (https://www.ebi.ac.uk/ena/browser/view/PRJEB62420) were converted to a HapMap using vcf2hapmap. The HapMap was filtered to keep only variants that were diallelic and was then split by breed using hapmapSampleIDsfromList. The hapmapGetROH tool was then used to extract regions of ROH per sample. ROH lengths were added together for each sample using sumCol, and the sums of ROH per sample were divided by the genome size to determine *F*_*ROH*_. *F*_*ROH*_ values for each breed’s HapMap were plotted in R (Fig. 8). These aligned with the observations by Meadows et al. (2023).

**Figure 8:**
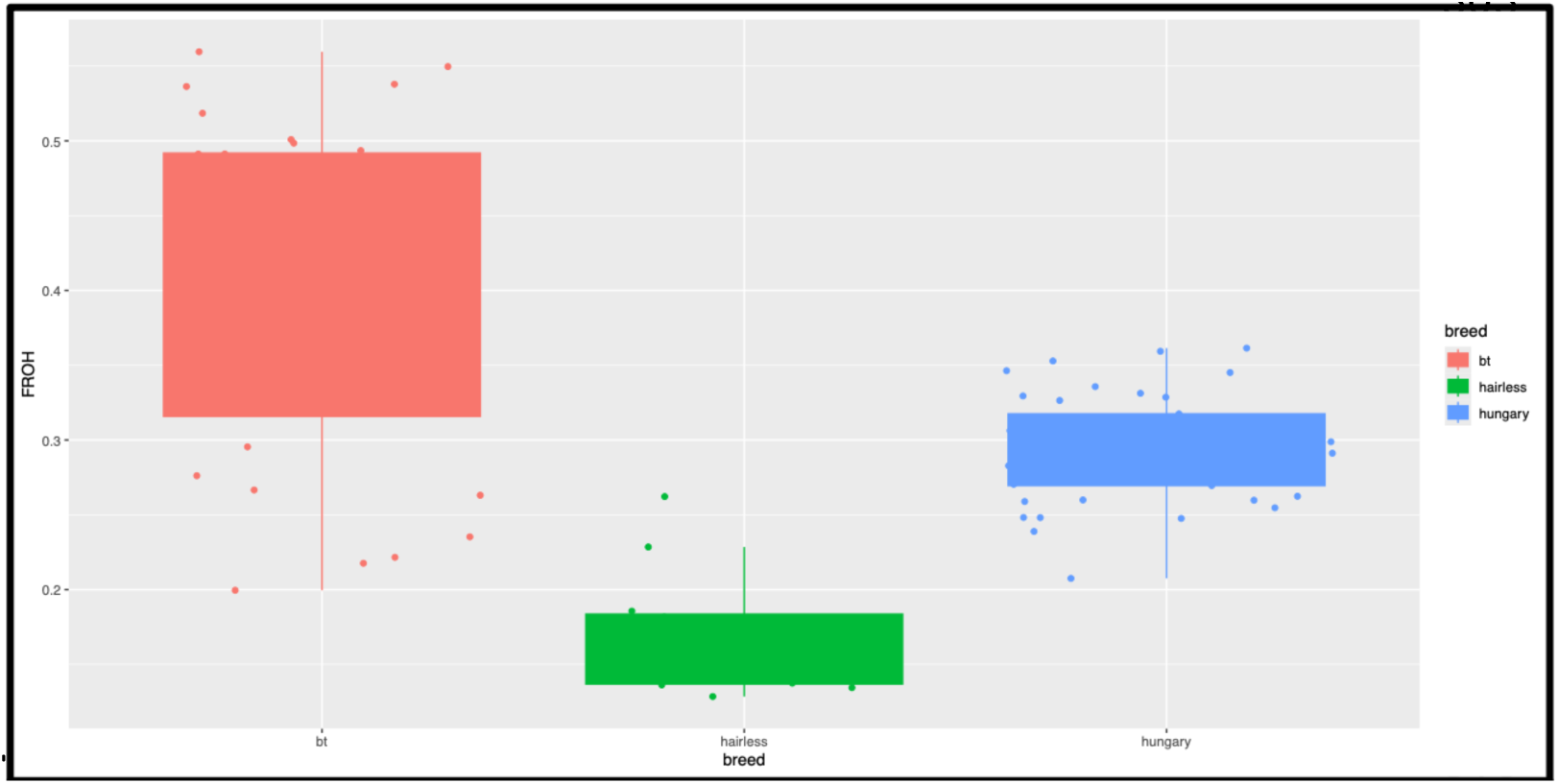
*F_ROH_* for three canine breeds using tools in the KhufuEnv. As was reported by Meadows et al. 2023, hairless dogs have relatively low *F_ROH_* while bt dogs have relatively higher *F_ROH_* and Hungary dogs have *F_ROH_* in between.

### Case study 4) Mapping QTL associated with hairlessness in canine

Canine ectodermal dysplasia (CED) is a condition commonly found in Mexican and Peruvian hairless dogs and Chinese crested dogs which results in hairlessness and abnormal tooth morphology (Drögemüller et al. 2008; Kupczik et al. 2017). CED is a semidominant, autosomal trait (Robinson 1985) for which heterozygous individuals exhibit the hairless phenotype while homozygotes are lethal. Previously, CED has been mapped to a region on chromosome 17 (Drögemüller et al. 2008; O’Brien et al. 2005), and further mutation and expression analyses supported the identification of the transcription factor *FOXI3* as a candidate gene for ectodermal development (Drögemüller et al. 2008).

Specifically, the mutant allele of *FOXI3* harbors a frameshift mutation likely rendering the protein nonfunctional (Drögemüller et al. 2008). Using the genotyping data published by Meadows et al. (2023) across 1600 dog samples including those which vary for the hairless trait, we attempted to recapitulate the candidate region associated with CED identified by Drögemüller et al. (2008) using the KhufuEnv tools (Fig. 9).

**Figure 9:**
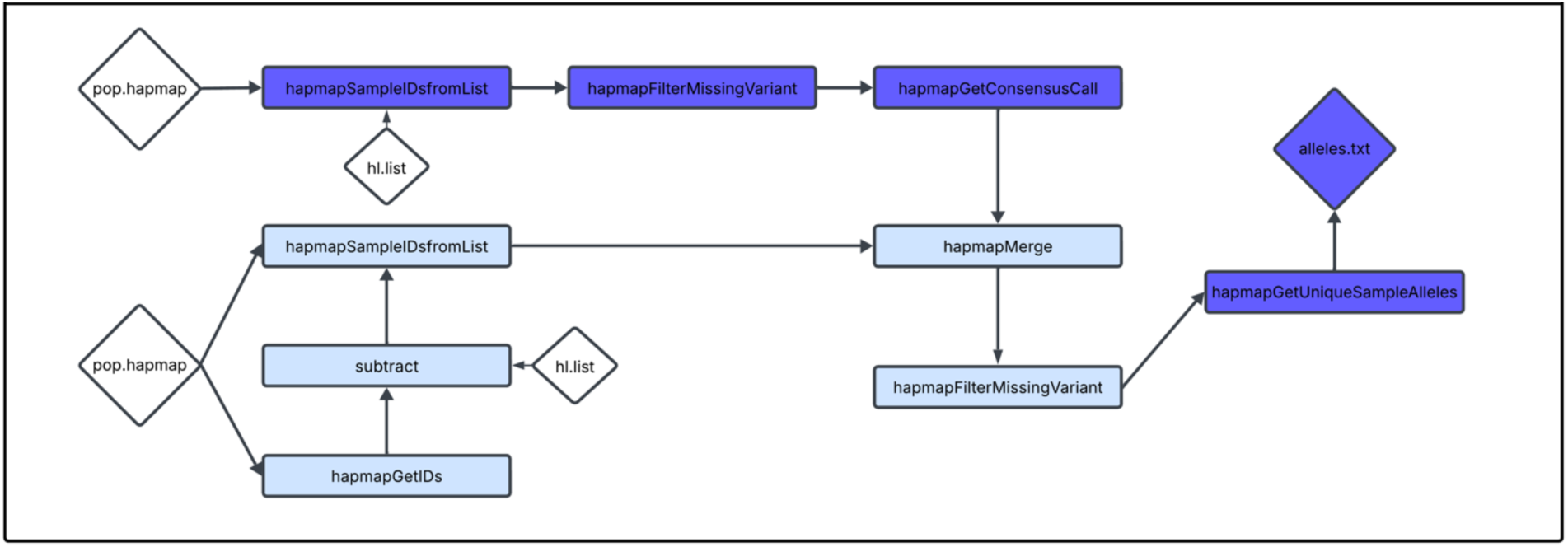
The custom KhufuEnv pipeline used for identifying QTL for CED in canine.

The previously generated HapMap for the variant calls by Meadows et al. (2023) was used to extract breed names with hapmapGetIDs. A list of non-hairless dog breeds was constructed by using the subtract tool to remove a list of the 11 hairless breeds from the initial sample list. Due to its large size, the initial VCF was split into 165 subsets (subsets of 100,000 variants) to allow for faster processing. The VCFs were converted to HapMaps with the vcf2hapmap tool. HapMaps corresponding to the hairless samples were generated with the hapmapSampleIDsfromList tool, supplied by the list of 11 hairless breeds. Variants were filtered to remove those with > 50% missing data, and the hapmapGetConsensusCall tool was used to extract the most common allele for each of the variants in the hairless samples. Then, HapMaps corresponding to the non-hairless samples were generated with the hapmapSampleIDsfromList tool using the non-hairless samples list. The non-hairless HapMaps were merged with the consensus hairless HapMap using hapmapMerge, and the merged HapMap was filtered to remove variants with > 50% missing data with hapmapFilterMissingVariant. Alleles unique to the hairless samples were extracted using hapmapGetUniqueSampleAlleles, and these were mined for candidate regions associated with CED. One region corresponded with the previously identified *FOXI3* gene associated with CED (Fig. 10).

**Figure 10:**
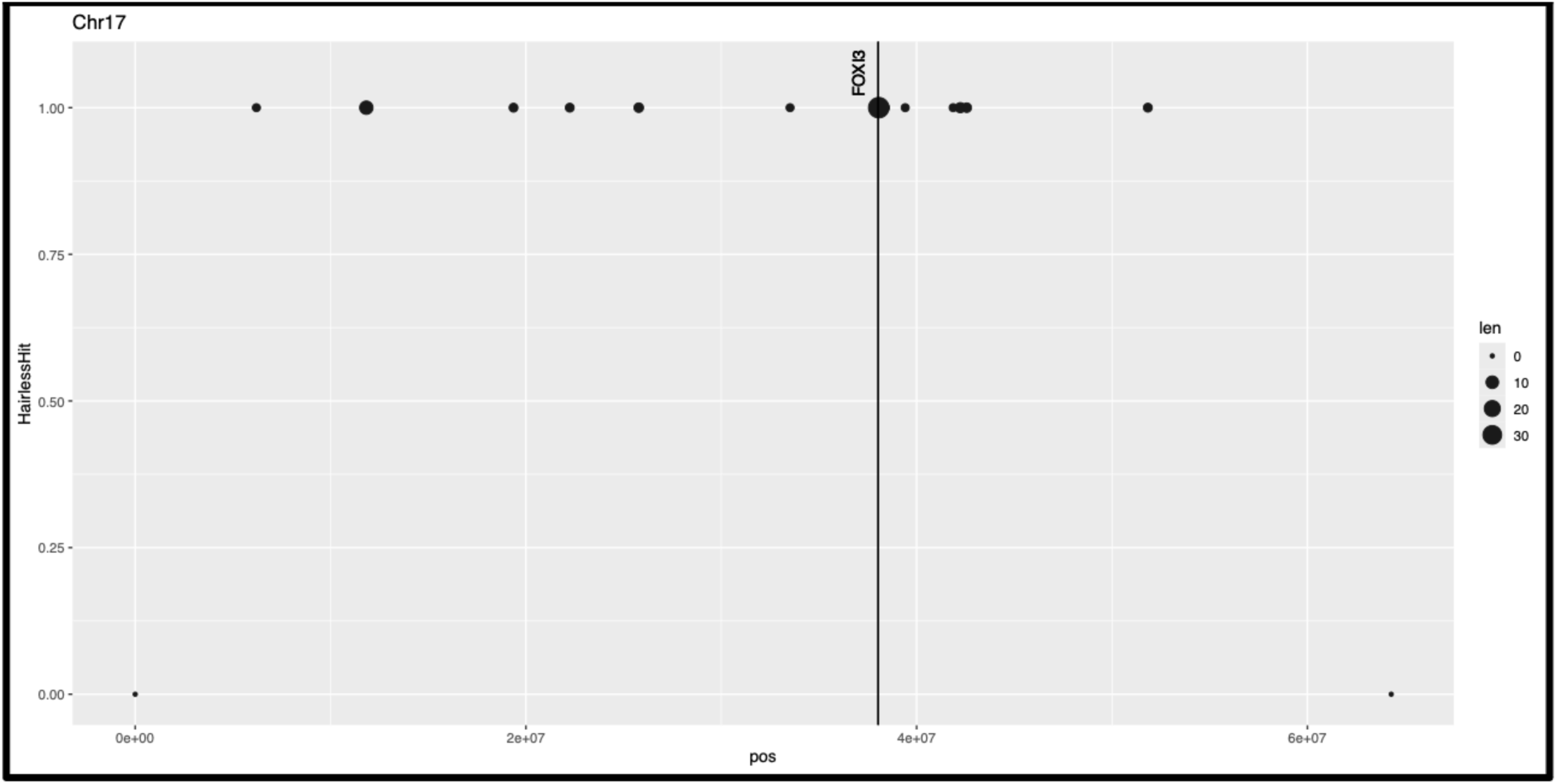
Identifying unique sample alleles between hairless and non-hairless dogs in the KhufuEnv identified a candidate region harboring a gene previously characterized by Drogemuller et al. 2008 for CED in canine. “Len” represents the maximum length of an allele variant at a respective location.

## Discussion

More genomics data is being generated than ever before. This year, it is expected that 40 exabytes will be needed to store genomics data (Stephens et al. 2015), equivalent to the data required to stream 1 billion hours (114,155 years) of standard definition content (Netflix 2025). This behemoth of data being generated necessitates the development and implementation of tools for rapid data analysis, in addition to expanding the genomics data science workforce (National Human Genome Research Institute 2021). To address these challenges, we developed the KhufuEnv, a user-friendly open-source auxiliary environment to assist analyzing genetics and genomics data. We have described some of the 132 tools present in its current release, along with the development of our custom HapMap and PanMap formats. While the standard HapMap has been a widely utilized file format for variant calls against linear genomes (The International HapMap Consortium 2003), to-date, there has yet to be an equivalent format developed for visualizing long stretches of SVs. By numerically coding allele calls, long SVs and shorter mutations can be visualized side-by-side to facilitate ease for data wrangling within Unix or GUI platforms.

In four case studies across animal and plant systems within autosome and allosome regions, we have benchmarked custom KhufuEnv pipelines to demonstrate the power and accuracy of its tools for trait mapping. Zhang et al. (2023) identified a 4.18 Mb region (152,644,698–156,822,195) region on chromosome 19 (B09) associated with cold tolerance during peanut germination, which was later fine-mapped to a 216 kb region (155,637,831–155,854,093) with Kompetitive Allele Specific PCR (KASP) markers. Beyond the variant calling, we accomplished finding a QTL near the fine-mapped region (155.2 Mb) (Fig. 4 and Supplementary File 1) using only nine lines of Unix code in a single environment, while the original authors required constructing a genetic map and QTL mapping with three softwares, SNPBinner (Gonda et al. 2019), Icimapping (Meng et al. 2015), and BLAT (Kent 2002), and two R packages, LinkageMapView (Ouellette et al. 2017) and R/qtl (Arends et al. 2010). Similarly, in the case of identifying the SDR region in *Amborella*, the original authors utilized an extensive approach that incorporated TRIMMOMATIC (Bolger et al. 2014) to generate k-mers for the WGS samples, identifying canonical 21-mers with Jellyfish (Marçais and Kingsford 2011), mapping all W-mers (female-specific k-mers) back to the haplotype assemblies with BWA-MEM, calculating coverage in sliding windows with BEDTools, and plotting with the R package karyoploteR (Gel and Serra 2017). We were able to accomplish identifying the SDR exclusively with KhufuEnv tools in seven lines of code beyond the initial pangenome graph construction and variant calling (Fig. 6A and 6B and Supplementary File 2).

Perhaps the greatest advantage of the KhufuEnv is its efficiency. In addition to exploiting tools to accurately *de novo* identify QTL which were previously characterized, even on a single thread, KhufuEnv tools identify such regions using at a fraction of the time for identifying regions as listed in the previous studies (Table 1). *De novo* identification of the cold tolerance and SDR regions took less than five minutes and three minutes, respectively. Calculating FROH across different breeds was done in less than six minutes, and the process of *de novo* identifying the hairless canine QTL too less than one hour to run all subsets. This marriage of the time efficiency and accuracy in *de novo* calling QTL in benchmarking datasets is perchance unprecedented.

**Table 1:** Processing time for different Khufu pipelines for benchmarking.

| Case Study | Reference | Maximum Khufu Processing Time |
| --- | --- | --- |
| Peanut cold tolerance | Zhanget al. 2023 | 4 min 0 sec |
| *Amborella* SDR | Carey et al. 2024 | 2 min 56 sec |
| Canine FROH | Meadows et al. 2023 | 2 min 20 sec to 6 min 0 sec |
| Canine hairlessness | Drögemüller et al. 2008 | 17 mins 22 sec to 40 mins 2 sec |

Other advantages the KhufuEnv presents are its ability read files from the standard input and write to standard output, allowing for more complex, pipeable actions to be done in a single line (with the exception of PanMap tools as PanMap tools output two files, a PanMap and a FASTA file). While Unix has several built-in tools that can be used for manipulating data files, some have limitations with regards to genomics data. For instance, “join” can be used to merge lines from two files based on a common field (Supplementary Fig. 4 and 5). When using join to merge two HapMap files, the join tool only merges data based on the first column, “chr” (Supplementary Fig. 6). In the KhufuEnv, however, we can use the hapmapMerge tool to combine two HapMaps by joining two input HapMaps in the order they are passed or the hapmapConcatenate tool to combine row data for shared sample ID’s in two HapMaps (Supplementary Fig. 7 and 8, respectively). Although general statistics tools are useful for obtaining global views on genotyping data (and other datasets), Unix does not include built-in statistics tools. The KhufuEnv statistics section can be extended to countless file types to easily extract statistics information.

While KhufuEnv offers an integrative set of tools to manipulate and analyze genomics data, we acknowledge its limitations. Zhang et al. (2023) developed a genetic map and exploited genetic linkage to identify QTL. Relying solely on phenotypic information and differences in allele frequencies between sample sets, we are not able to determine the phenotypic variation explained (PVE), *R*^2^, or if the allele of interest behaves in an additive, dominant, or epistatic fashion. When utilizing our KhufuEnv approach for identifying a QTL for CED in canine (Fig. 10), it is relevant to note in this approach that several significant hits were found across the genome outside the region harboring *FOXI3* (Supplementary File 3). However, this is likely an artifact due to the small number of hairless dogs (11) we had to compare to a diverse group of over 1,000 non-hairless dogs. The study by Drögemüller et al. (2008) utilized data for 20 hairless and 19 coated Chinese Crested dogs for their genome-wide association study (GWAS). Having nearly double the number of hairless individuals than our study and closer related individuals without the trait of interest to compare with likely resolves the noise experienced in our benchmark.

The hapmapGetROH tool in the current KhufuEnv release classifies ROH with a default stretch of 50 minimum bp of homozygosity. While this minimum can be user-customized, we have not yet incorporated a sliding window approach or customizations for allowable heterozygous sites within a window but anticipate improving this tool accordingly for the next release. Nonetheless, our results mimic the pattern demonstrated by Meadows et al. (2023) (Fig. 8 and Supplementary File 4).

In its current state, the KhufuEnv prioritizes simple and quick processes which can be run on a single thread. We recognize that this prevents the integration other tools such as DNA sequence aligners like BWA (Li and Durbin 2009) or variant callers like GATK (McKenna et al. 2010) into the environment itself and that users may have to rely on these programs during initial analyses if variant call files are not publicly available to them. Our needs for trait mapping inspired the present generation of KhufuEnv tools. The KhufuEnv currently does not have tools explicitly designed for manipulating GFF, BED, BAM, or SAM files (although some tools in our data set processing section may be applicable) for which other suites, such as BEDTools and BBTools, support. As KhufuEnv is an open-source resource, we will be integrating user feedback to design new tools dependent on community needs. Thus, we anticipate developing tools to specifically address other file types and data processing in future releases.

## Conclusion

We report the inaugural release of the KhufuEnv, an open-source toolbox to support genomic analysis. The KhufuEnv allows users to develop modular, custom pipelines for analyzing and manipulating genomics data in a standalone environment, and the custom HapMap and PanMap formats allow for visualizing variants called from linear and pangenomes, respectively, in a concise manner. As demonstrated by four case studies pertaining to plant and animal breeding, KhufuEnv tools can facilitate accurate and rapid *de novo* identification of relevant QTL for animal and plant breeding. While its current state prioritizes speed, simplicity, and single-threaded tools, we look forward to incorporating additional tools and capabilities to suit user preferences in future releases.

## Availability of Supporting Source Code and Requirements

KhufuEnv tools are compatible with Unix systems. R and gawk are required dependencies for the KhufuEnv.

## Example Code

As an example for the custom pipelines we have built with KhufuEnv tools, we have provided the hard code for the cold tolerance peanut benchmark study which corresponds to Figure 3:

cat par_min5.hapmap | hapmapSNP2SV | hapmapGetDiAlleles | hapmapGetHomoPolymorphic > par.hapmap

hapmapMerge par.hapmap pop_min3.hapmap | hapmapSNP2SV | hapmapGetDiAlleles | hapmapFilterMissingVariant - 0.5 > pop.hapmap

for i in $(seq 2 6); do

cat data.txt | cut -f 1,$i | sort -k2,2nr | cut -f 1| head -20 | sed “1iHuayu44” > high.list; cat data.txt | cut -f 1,$i | sort -k2,2n | cut -f 1| head -20 | sed “1iHuayu44” > low.list hapmapSampleIDsfromList pop.hapmap high.list | hapmapGetFreqSampleAlleles - “Huayu44” | awk ’{if($5>=4) print $1”_”$2”\t”$4}’ > highpop.freq hapmapSampleIDsfromList pop.hapmap low.list | hapmapGetFreqSampleAlleles - “Huayu44” | awk ’{if($5>=4) print $1“_”$2“\t”$4}’ > lowpop.freq

merge highpop.freq lowpop.freq | grep -v NA | sed 1d | awk ’{x=$2-$3; printf “%s\t%0.3f\n”,$1, sqrt(x*x)}’ | sed “s:_:\t:g” | sed “s:$:\t$((i-1)):g”

done | sed “1ichr\tpos\tdeltaSNP\tenv” > freq.txt

## Funding

This work was supported by NIFA award 2023-78408-39694 and a strategic partnership with the city of Dothan, AL.

## Conflict of Interest

Josh Clevenger is a co-founder of Veil Genomics.

## Acknowledgements

Thank you to Richard Johnson and the IT team at HudsonAlpha for supporting the HPC resources utilized for this work.

## Author Contributions

W. K. and J. C. conceived the idea of the KhufuEnv and benchmarking case studies, H. W. wrote the manuscript, W. K. and H. W. generated figures, W. K. wrote the tools for the environment, H. W. and C. D. troubleshot and contributed to tool development, C. D. wrote documentation, J. C. oversaw the experimental design and provided associated funding, and all authors contributed to and approved the final draft.

## Data Availability

The KhufuEnv can be downloaded at https://github.com/w-korani/KhufuEnv. All raw data used in this manuscript was derived from prior experiments associated with their respective citations. Testing datasets are available within the KhufuEnv. HapMaps and PanMaps generated from the Khufu platform are available upon request.

## Supplementary Figures

**Supplementary Figure 1:**
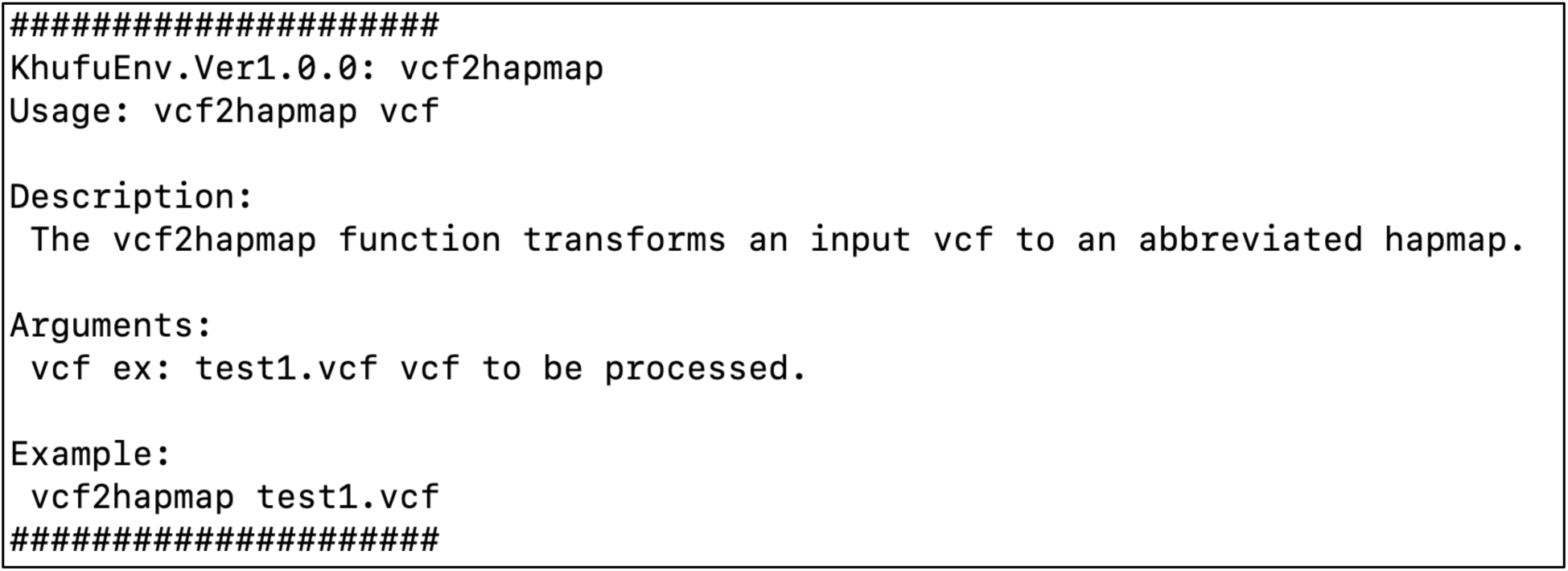
The KhufuEnvHelp [tool] command outputs information regarding a tool’s description, parameters, and practice files for using the tool. This is an example output from running the command “KhufuEnvHelp vcf2hapmap”.

**Supplementary Figure 2:**
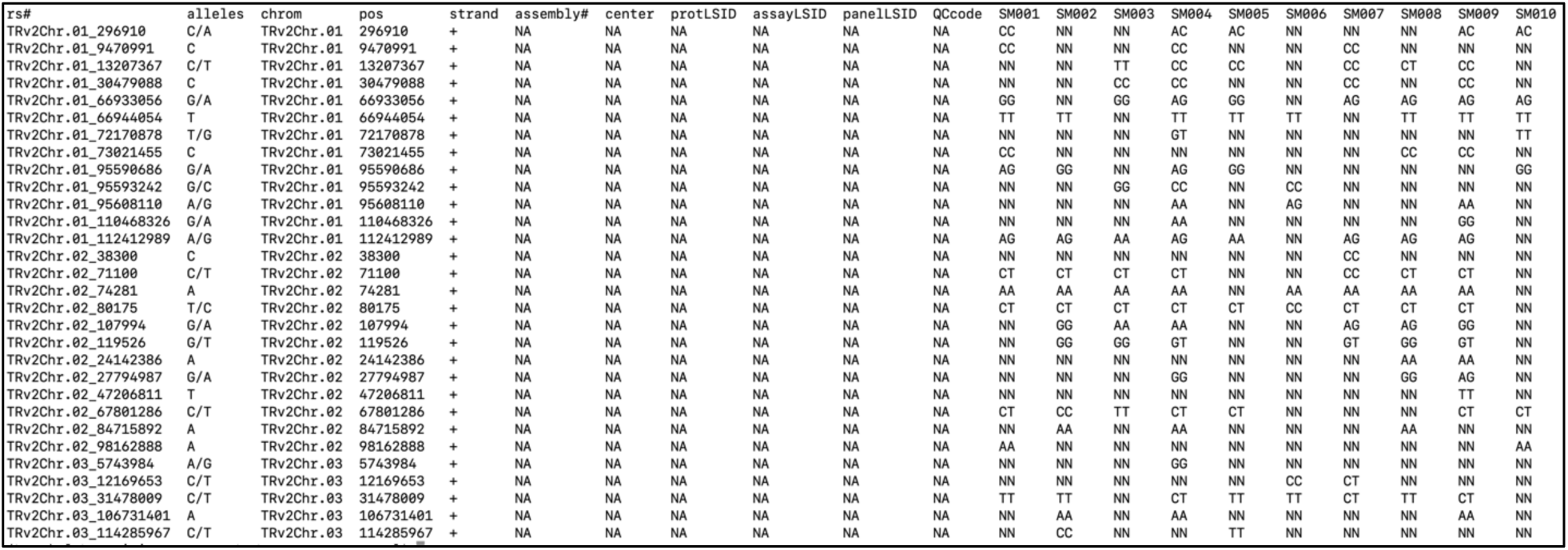
An example of the standard HapMap file format developed by The International HapMap Consortium (2003). This is an example output from running the command “cat test1.stdhapmap | column -t”.

**Supplementary Figure 3:**
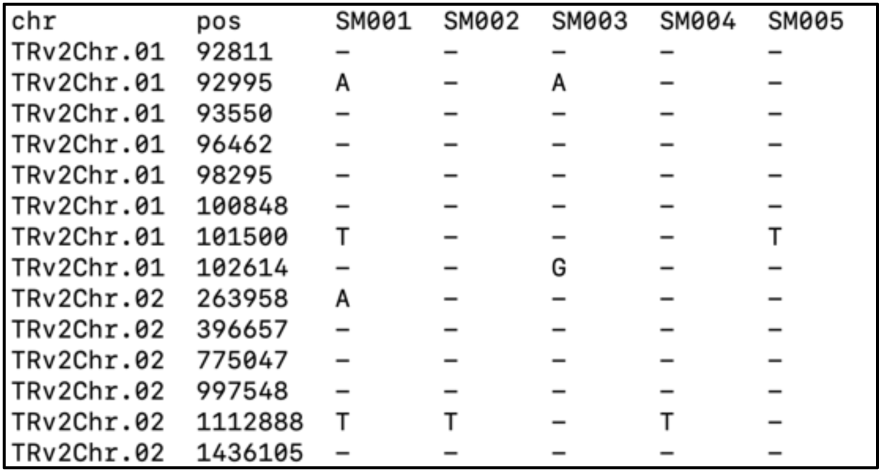
An example of the condense Khufu HapMap format. This is an example output from converting the test1.stdhapmap to a Khufu HapMap running the command “stdhapmap2hapmap test1.stdhapmap > test1.std.toHAPMAP” then “cat test1.std.toHAPMAP | column -t”.

**Supplementary Figure 4:**
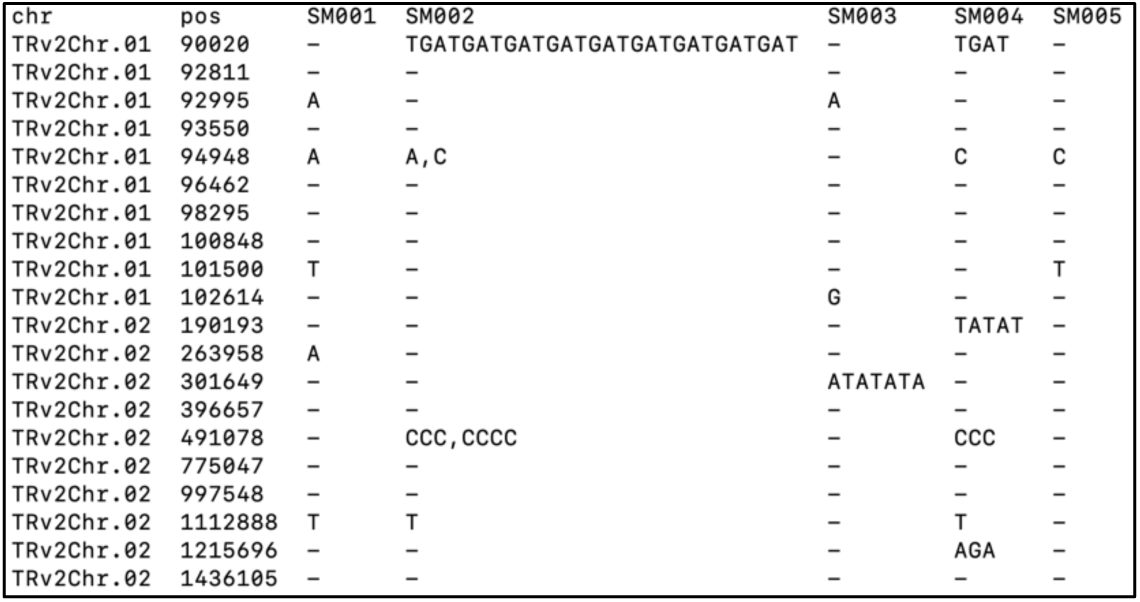
An example HapMap displayed by running the command “cat test1.hapmap | column -t”.

**Supplementary Figure 5:**
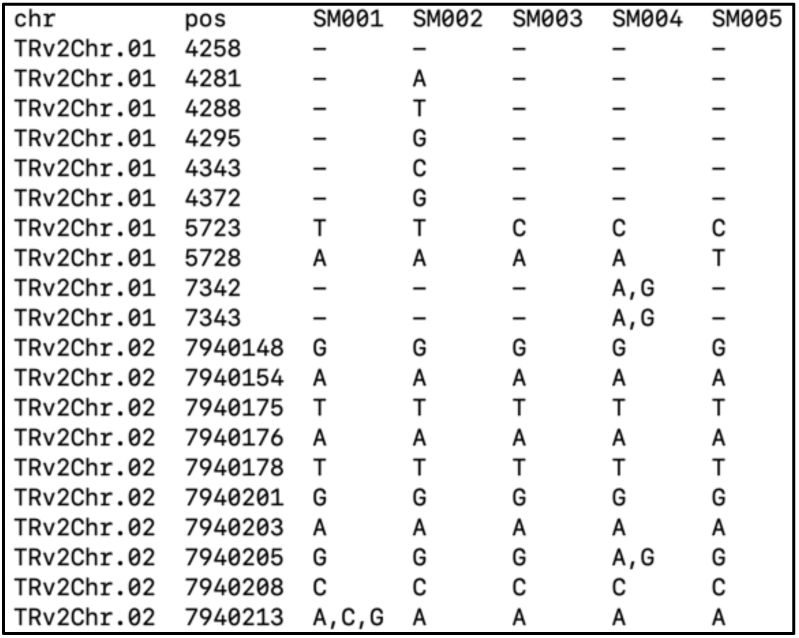
An example HapMap displayed by running the command “cat test2.hapmap | column -t”.

**Supplementary Figure 6:**
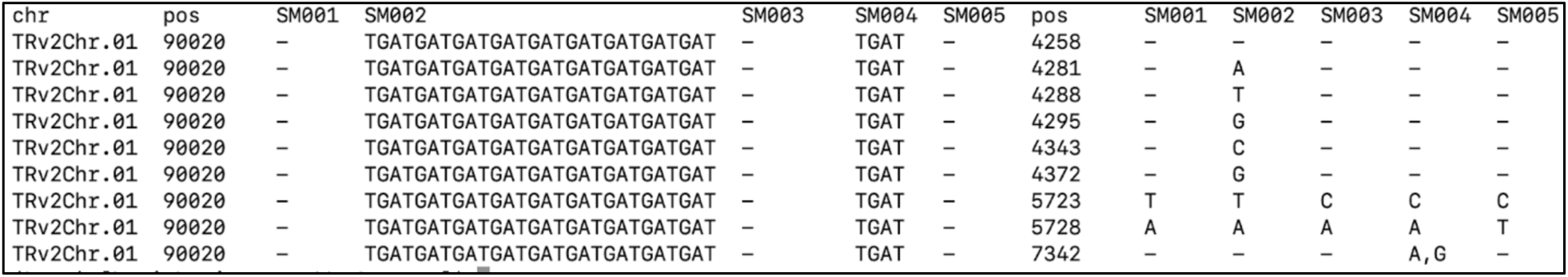
Instead of combining the genotyping data for each of the samples from the HapMaps in Supplementary Figs. 4 and 5, the join function in Unix only merges data based on the first column when using the commands “merge test1.hapmap test2.hapmap > merged.hapmap” and “cat merged.hapmap | column -t | head”.

**Supplementary Figure 7:**
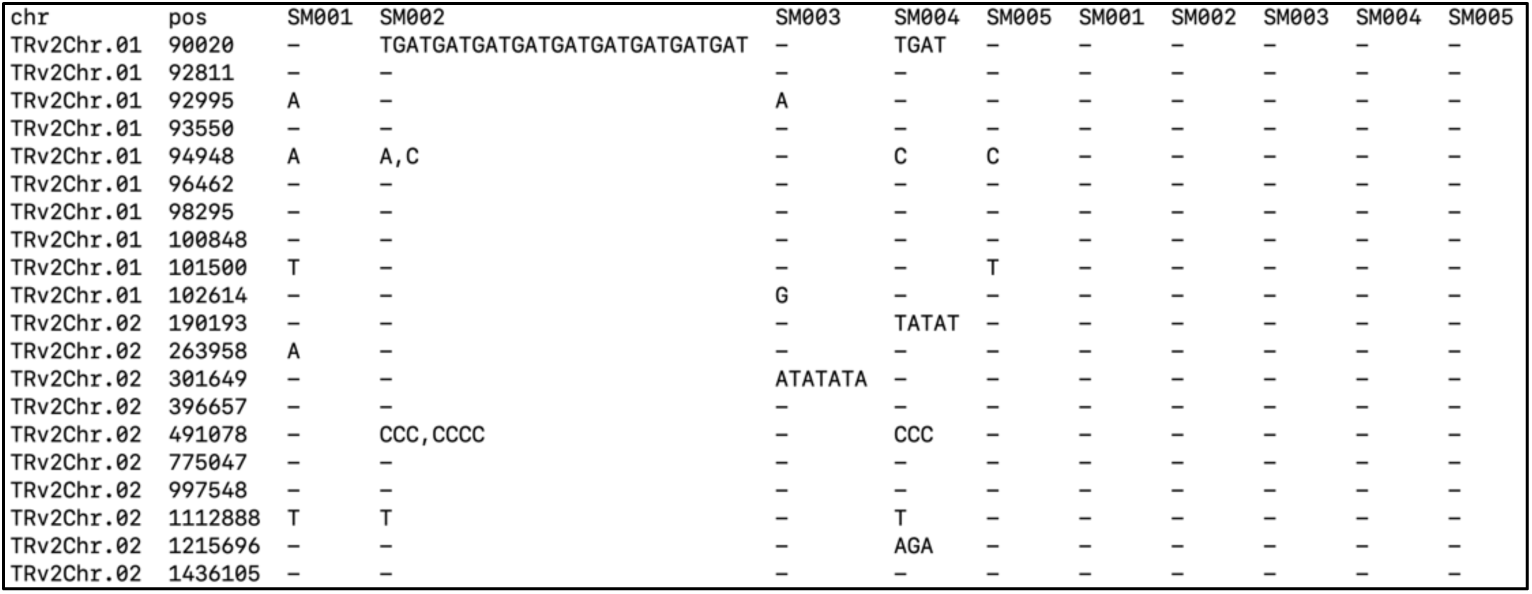
Alternative to the Unix join function, the KhufuEnv hapmapMerge tool combines information from two HapMaps such that row data existing in the right (secondary) HapMap that does not exist in the left HapMap (primary) will be dropped. Commands used to generate the merged HapMap were “hapmapMerge test1.hapmap test2.hapmap > Merge.hapmap” and “cat Merge.hapmap | column -t”.

**Supplementary Figure 8:**
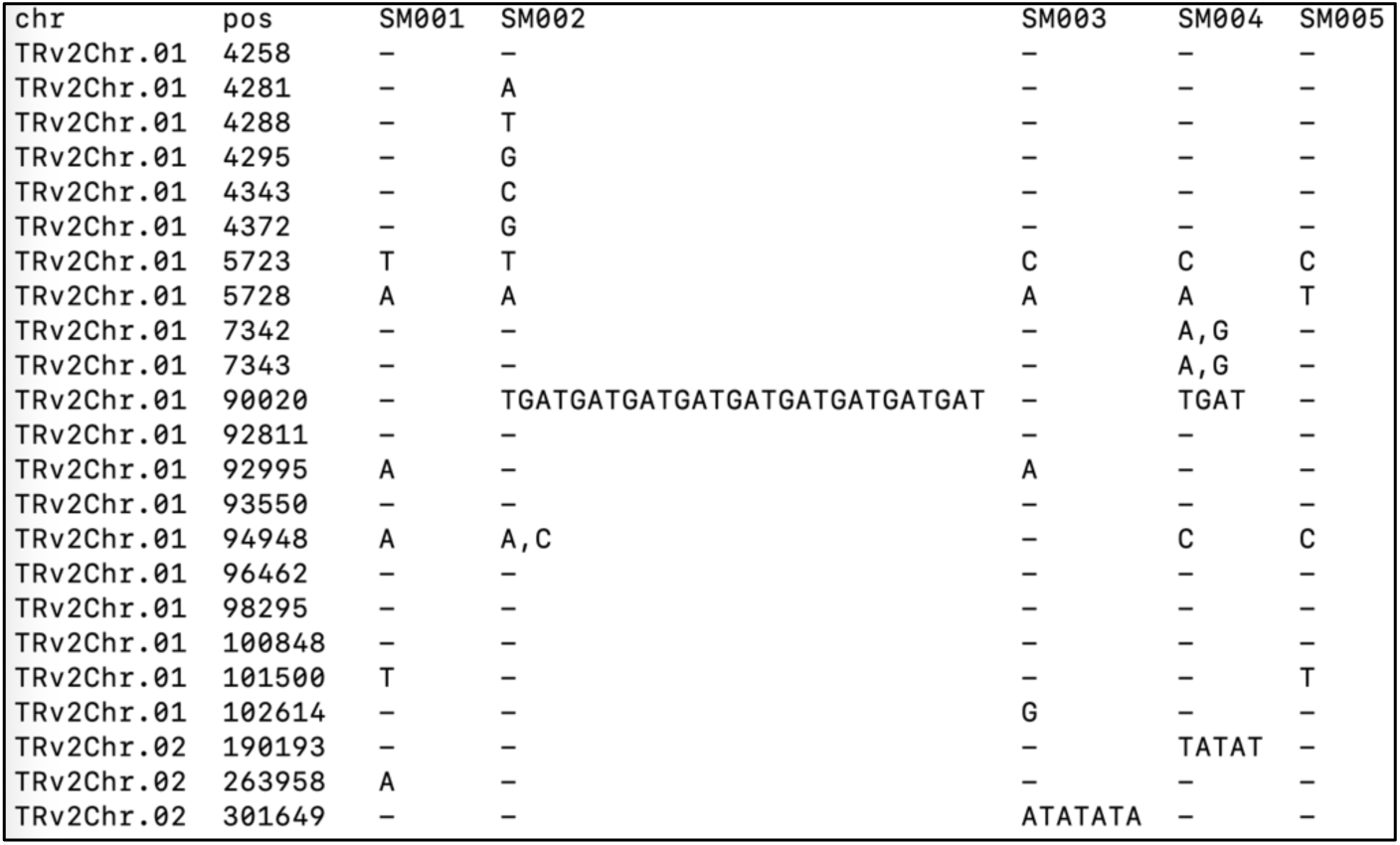
The KhufuEnv hapmapConcatenate tool combines row data for samples of shared ID’s in two input HapMaps. Commands used to generate the merged HapMap were “hapmapConcatenate test1.hapmap test2.hapmap > Concat.hapmap” and “cat Concat.hapmap | column -t”.

## References

1. Adams K (2013) Genomic Clues to the Ancestral Flowering Plant. Science 342:1456–1457

2. Anger N, Fogliani B, Scutt CP, Gâteblé G (2017) Dioecy in *Amborella trichopoda*: evidence for genetically based sex determination and its consequences for inferences of the breeding system in early angiosperms. Annals of Botany 119:591- 597

3. Arends D, Prins P, Jansen RC, Broman KW (2010) R/qtl: high-throughput multiple QTL mapping. Bioinformatics 26:2990–2992

4. Bartlett C (2014) The Design of The Great Pyramid of Khufu. Nexus Network Journal 16:299–311

5. Bayat A (2002) Science, medicine, and the future: Bioinformatics. Bmj 324:1018–1022

6. Bolger AM, Lohse M, Usadel B (2014) Trimmomatic: a flexible trimmer for Illumina sequence data. Bioinformatics 30:2114-2120

7. Broman KW, Weber JL (1999) Long homozygous chromosomal segments in reference families from the centre d’Etude du polymorphisme humain. The American Journal of Human Genetics 65:1493–1500

8. Bushnell B, Rood J, Singer E (2017) BBMerge – Accurate paired shotgun read merging via overlap. PLOS ONE 12:e0185056

9. Carey SB, Aközbek L, Lovell JT, Jenkins J, Healey AL, Shu S, Grabowski P, Yocca A, Stewart A, Jones T, Barry K, Rajasekar S, Talag J, Scutt C, Lowry PP, Munzinger J, Knox EB, Soltis DE, Soltis PS, Grimwood J, Schmutz J, Leebens-Mack J, Harkess A (2024) ZW sex chromosome structure in *Amborella trichopoda*. Nature Plants 10:1944–1954

10. Charlesworth D (2002) Plant sex determination and sex chromosomes. Heredity 88:94–101

11. Charlesworth D, Harkess A (2024) Why should we study plant sex chromosomes? The Plant Cell 36:1242–1256

12. Chu Y, Chee P, Culbreath A, Isleib TG, Holbrook CC, Ozias-Akins P (2019) Major QTLs for Resistance to Early and Late Leaf Spot Diseases Are Identified on Chromosomes 3 and 5 in Peanut (*Arachis hypogaea*). Frontiers in Plant Science Volume 10 – 2019

13. Clevenger J, Chu Y, Chavarro C, Botton S, Culbreath A, Isleib TG, Holbrook CC, Ozias-Akins P (2018) Mapping Late Leaf Spot Resistance in Peanut (*Arachis hypogaea*) Using QTL-seq Reveals Markers for Marker-Assisted Selection. Frontiers in Plant Science Volume 9 – 2018

14. The International HapMap Consortium (2003) The International HapMap Project. Nature 426:789-796

15. Crossa J, Campos Gde L, Pérez P, Gianola D, Burgueño J, Araus JL, Makumbi D, Singh RP, Dreisigacker S, Yan J, Arief V, Banziger M, Braun HJ (2010) Prediction of genetic values of quantitative traits in plant breeding using pedigree and molecular markers. Genetics 186:713–724

16. Danecek P, Auton A, Abecasis G, Albers CA, Banks E, DePristo MA, Handsaker RE, Lunter G, Marth GT, Sherry ST, McVean G, Durbin R (2011) The variant call format and VCFtools. Bioinformatics 27:2156–2158

17. Das S, Brown J (2024) Oilseeds and Products Update. United States Department of Agriculture Foreign Agricultural Service

18. Drögemüller C, Karlsson EK, Hytönen MK, Perloski M, Dolf G, Sainio K, Lohi H, Lindblad-Toh K, Leeb T (2008) A mutation in hairless dogs implicates *FOXI3* in ectodermal development. Science 321:1462

19. Ferenčaković M, Hamzić E, Gredler B, Solberg TR, Klemetsdal G, Curik I, Sölkner J (2013) Estimates of autozygosity derived from runs of homozygosity: empirical evidence from selected cattle populations. Journal of Animal Breeding and Genetics 130:286–293

20. Gel B, Serra E (2017) karyoploteR: an R/Bioconductor package to plot customizable genomes displaying arbitrary data. Bioinformatics 33:3088–3090

21. Gonda I, Ashrafi H, Lyon DA, Strickler SR, Hulse-Kemp AM, Ma Q, Sun H, Stoffel K, Powell AF, Futrell S, Thannhauser TW, Fei Z, Van Deynze AE, Mueller LA, Giovannoni JJ, Foolad MR (2019) Sequencing-Based Bin Map Construction of a Tomato Mapping Population, Facilitating High-Resolution Quantitative Trait Loci Detection. The Plant Genome 12:180010

22. Gorssen W, Meyermans R, Janssens S, Buys N (2021) A publicly available repository of ROH islands reveals signatures of selection in different livestock and pet species. Genetics Selection Evolution 53:2

23. Hickey G, Monlong J, Ebler J, Novak AM, Eizenga JM, Gao Y, Abel HJ, Antonacci-Fulton LL, Asri M, Baid G, Baker CA, Belyaeva A, Billis K, Bourque G, Buonaiuto S, Carroll A, Chaisson MJP, Chang P-C, Chang XH, Cheng H, Chu J, Cody S, Colonna V, Cook DE, Cook-Deegan RM, Cornejo OE, Diekhans M, Doerr D, Ebert P, Ebler J, Eichler EE, Fairley S, Fedrigo O, Felsenfeld AL, Feng X, Fischer C, Flicek P, Formenti G, Frankish A, Fulton RS, Garg S, Garrison E, Garrison NA, Giron CG, Green RE, Groza C, Guarracino A, Haggerty L, Hall IM, Harvey WT, Haukness M, Haussler D, Heumos S, Hoekzema K, Hourlier T, Howe K, Jain M, Jarvis ED, Ji HP, Kenny EE, Koenig BA, Kolesnikov A, Korbel JO, Kordosky J, Koren S, Lee H, Lewis AP, Liao W-W, Lu S, Lu T- Y, Lucas JK, Magalhães H, Marco-Sola S, Marijon P, Markello C, Marschall T, Martin FJ, McCartney A, McDaniel J, Miga KH, Mitchell MW, Mountcastle J, Munson KM, Mwaniki MN, Nattestad M, Nurk S, Olsen HE, Olson ND, Pesout T, Phillippy AM, Popejoy AB, Porubsky D, Prins P, Puiu D, Rautiainen M, Regier AA, Rhie A, Sacco S, Sanders AD, Schneider VA, Schultz BI, Shafin K, Sibbesen JA, Sirén J, Smith MW, Sofia HJ, Tayoun ANA, Thibaud-Nissen F, Tomlinson C, Tricomi FF, Villani F, Vollger MR, Wagner J, Walenz B, Wang T, Wood JMD, Zimin AV, Zook JM, Marschall T, Li H, Paten B, Human Pangenome Reference C (2024) Pangenome graph construction from genome alignments with Minigraph-Cactus. Nature Biotechnology 42:663–673

24. Hurdle NL, Grey TL, Pilon C, Monfort WS, Prostko EP (2020) Peanut Seed Germination and Radicle Development Response to Direct Exposure of Flumioxazin Across Multiple Temperatures. Peanut Science 47:89–93

25. Kent WJ (2002) BLAT--the BLAST-like alignment tool. Genome Research 12:656–664

26. Kim J, Macharia JK, Kim M, Heo JM, Yu M, Choo HJ, Lee JH (2024) Runs of homozygosity analysis for selection signatures in the Yellow Korean native chicken. Animal Bioscience 37:1683–1691

27. Korani W, O’Connor D, Chu Y, Chavarro C, Ballen C, Guo B, Ozias-Akins P, Wright G, Clevenger J (2021) De novo QTL-seq Identifies Loci Linked to Blanchability in Peanut (*Arachis hypogaea*) and Refines Previously Identified QTL with Low Coverage Sequence. Agronomy 11:2201

28. Kupczik K, Cagan A, Brauer S, Fischer MS (2017) The dental phenotype of hairless dogs with *FOXI3* haploinsufficiency. Scientific Reports 7:5459

29. Li H (2013)Seqtk.

30. Li H (2017) bioawk.

31. Li H, Durbin R (2009) Fast and accurate short read alignment with Burrows–Wheeler transform. Bioinformatics 25:1754–1760

32. Marçais G, Kingsford C (2011) A fast, lock-free approach for efficient parallel counting of occurrences of k-mers. Bioinformatics 27:764–770

33. Mastrangelo S, Sardina MT, Tolone M, Di Gerlando R, Sutera AM, Fontanesi L, Portolano B (2018) Genome-wide identification of runs of homozygosity islands and associated genes in local dairy cattle breeds. Animal 12:2480–2488

34. McKenna A, Hanna M, Banks E, Sivachenko A, Cibulskis K, Kernytsky A, Garimella K, Altshuler D, Gabriel S, Daly M, DePristo MA (2010) The Genome Analysis Toolkit: a MapReduce framework for analyzing next-generation DNA sequencing data. Genome Research 20:1297–1303

35. Meadows JRS, Kidd JM, Wang G-D, Parker HG, Schall PZ, Bianchi M, Christmas MJ, Bougiouri K, Buckley RM, Hitte C, Nguyen AK, Wang C, Jagannathan V, Niskanen JE, Frantz LAF, Arumilli M, Hundi S, Lindblad-Toh K, Ginja C, Agustina KK, André C, Boyko AR, Davis BW, Drögemüller M, Feng X-Y, Gkagkavouzis K, Iliopoulos G, Harris AC, Hytönen MK, Kalthoff DC, Liu Y-H, Lymberakis P, Poulakakis N, Pires AE, Racimo F, Ramos-Almodovar F, Savolainen P, Venetsani S, Tammen I, Triantafyllidis A, vonHoldt B, Wayne RK, Larson G, Nicholas FW, Lohi H, Leeb T, Zhang Y-P, Ostrander EA (2023) Genome sequencing of 2000 canids by the Dog10K consortium advances the understanding of demography, genome function and architecture. Genome Biology 24:187

36. Meng L, Li H, Zhang L, Wang J (2015) QTL IciMapping: Integrated software for genetic linkage map construction and quantitative trait locus mapping in biparental populations. The Crop Journal 3:269–283

37. Meuwissen THE, Hayes BJ, Goddard ME (2001) Prediction of Total Genetic Value Using Genome-Wide Dense Marker Maps. Genetics 157:1819–1829

38. Meyermans R, Gorssen W, Buys N, Janssens S (2020) How to study runs of homozygosity using PLINK? A guide for analyzing medium density SNP data in livestock and pet species. BMC Genomics 21:94

39. Mulim HA, Brito LF, Pinto LFB, Ferraz JBS, Grigoletto L, Silva MR, Pedrosa VB (2022) Characterization of runs of homozygosity, heterozygosity-enriched regions, and population structure in cattle populations selected for different breeding goals. BMC Genomics 23:209

40. National Human Genome Research Institute (2021) NHGRI Action Agenda for Genomics Workforce Diversity. National Institute for Health

41. NCBI (2020) We want to hear from you about changes to NIH’s Sequence Read Archive data format and storage. National Center for Biotechnology Information

42. Netflix (2025) How to control how much data Netflix uses.

43. O’Brien DP, Johnson GS, Schnabel RD, Khan S, Coates JR, Johnson GC, Taylor JF (2005) Genetic mapping of canine multiple system degeneration and ectodermal dysplasia loci. Journal of Heredity 96:727–734

44. Ouellette LA, Reid RW, Blanchard SG, Brouwer CR (2017) LinkageMapView—rendering high-resolution linkage and QTL maps. Bioinformatics 34:306–307

45. Peripolli E, Munari DP, Silva MVGB, Lima ALF, Irgang R, Baldi F (2017) Runs of homozygosity: current knowledge and applications in livestock. Animal Genetics 48:255–271

46. Purcell S, Neale B, Todd-Brown K, Thomas L, Ferreira MA, Bender D, Maller J, Sklar P, de Bakker PI, Daly MJ, Sham PC (2007) PLINK: a tool set for whole-genome association and population-based linkage analyses. American Journal of Human Genetics 81:559–575

47. Quinlan AR, Hall IM (2010) BEDTools: a flexible suite of utilities for comparing genomic features. Bioinformatics 26:841–842

48. R Core Team (2021) R: A language and environment for statistical computing. R Foundation for Statistical Computing, Vienna, Austria

49. Renner SS (2014) The relative and absolute frequencies of angiosperm sexual systems: Dioecy, monoecy, gynodioecy, and an updated online database. American Journal of Botany 101:1588–1596

50. Robinson R (1985) Chinese crested dog. Journal of Heredity 76:217–218

51. Sams AJ, Boyko AR (2019) Fine-Scale Resolution of Runs of Homozygosity Reveal Patterns of Inbreeding and Substantial Overlap with Recessive Disease Genotypes in Domestic Dogs. G3 (Bethesda) 9:117-123

52. Shen W, Le S, Li Y, Hu F (2016) SeqKit: A Cross-Platform and Ultrafast Toolkit for FASTA/Q File Manipulation. PLOS ONE 11:e0163962

53. Shen W, Sipos B, Zhao L (2024) SeqKit2: A Swiss army knife for sequence and alignment processing. iMeta 3:e191

54. Sirén J, Monlong J, Chang X, Novak AM, Eizenga JM, Markello C, Sibbesen JA, Hickey G, Chang PC, Carroll A, Gupta N, Gabriel S, Blackwell TW, Ratan A, Taylor KD, Rich SS, Rotter JI, Haussler D, Garrison E, Paten B (2021) Pangenomics enables genotyping of known structural variants in 5202 diverse genomes. Science 374:abg8871

55. Soltis DE, Smith SA, Cellinese N, Wurdack KJ, Tank DC, Brockington SF, Refulio-Rodriguez NF, Walker JB, Moore MJ, Carlsward BS, Bell CD, Latvis M, Crawley S, Black C, Diouf D, Xi Z, Rushworth CA, Gitzendanner MA, Sytsma KJ, Qiu Y-L, Hilu KW, Davis CC, Sanderson MJ, Beaman RS, Olmstead RG, Judd WS, Donoghue MJ, Soltis PS (2011) Angiosperm phylogeny: 17 genes, 640 taxa. American Journal of Botany 98:704-730

56. Stephens ZD, Lee SY, Faghri F, Campbell RH, Zhai C, Efron MJ, Iyer R, Schatz MC, Sinha S, Robinson GE (2015) Big Data: Astronomical or Genomical? PLoS Biology 13:e1002195

57. Subramanian S, Kumar M (2024) The Association between the Abundance of Homozygous Deleterious Variants and the Morbidity of Dog Breeds. Biology 13:574

58. Telatin A, Fariselli P, Birolo G (2021) SeqFu: A Suite of Utilities for the Robust and Reproducible Manipulation of Sequence Files. Bioengineering 8

59. The International HapMap Consortium (2003) The International HapMap Project. Nature 426:789-796

60. Thorisson GA, Smith AV, Krishnan L, Stein LD (2005) The International HapMap Project Web site. Genome Research 15:1592–1593

61. Variath MT, Janila P (2017) Economic and Academic Importance of Peanut. In: Varshney RK, Pandey MK, Puppala N (eds) The Peanut Genome. Springer International Publishing, Cham, pp 7-26

62. Wang C, Wallerman O, Arendt M-L, Sundström E, Karlsson Å, Nordin J, Mäkeläinen S, Pielberg GR, Hanson J, Ohlsson Å, Saellström S, Rönnberg H, Ljungvall I, Häggström J, Bergström TF, Hedhammar Å, Meadows JRS, Lindblad-Toh K (2021) A novel canine reference genome resolves genomic architecture and uncovers transcript complexity. Communications Biology 4:185

63. Wiggans GR, Cole JB, Hubbard SM, Sonstegard TS (2017) Genomic Selection in Dairy Cattle: The USDA Experience. Annual Review of Animal Biosciences 5:309–327

64. Zhang H, Dong J, Zhao X, Zhang Y, Ren J, Xing L, Jiang C, Wang X, Wang J, Zhao S, Yu H (2019) Research Progress in Membrane Lipid Metabolism and Molecular Mechanism in Peanut Cold Tolerance. Frontiers in Plant Science 10

65. Zhang Q, Guldbrandtsen B, Bosse M, Lund MS, Sahana G (2015) Runs of homozygosity and distribution of functional variants in the cattle genome. BMC Genomics 16:542

66. Zhang X, Zhang X, Wang L, Liu Q, Liang Y, Zhang J, Xue Y, Tian Y, Zhang H, Li N, Sheng C, Nie P, Feng S, Liao B, Bai D (2023) Fine mapping of a QTL and identification of candidate genes associated with cold tolerance during germination in peanut (*Arachis hypogaea* L.) on chromosome B09 using whole genome re-sequencing. Frontiers in Plant Science 14:1153293

67. Carey SB, Aközbek L, Lovell JT, Jenkins J, Healey AL, Shu S, Grabowski P, Yocca A, Stewart A, Jones T, Barry K, Rajasekar S, Talag J, Scutt C, Lowry PP, Munzinger J, Knox EB, Soltis DE, Soltis PS, Grimwood J, Schmutz J, Leebens-Mack J, Harkess A (2024) ZW sex chromosome structure in *Amborella trichopoda*. Nature Plants 10:1944–1954

68. Kent WJ (2002) BLAT--the BLAST-like alignment tool. Genome Research 12:656–664

69. Korani W, O’Connor D, Chu Y, Chavarro C, Ballen C, Guo B, Ozias-Akins P, Wright G, Clevenger J (2021) De novo QTL-seq Identifies Loci Linked to Blanchability in Peanut (*Arachis hypogaea*) and Refines Previously Identified QTL with Low Coverage Sequence. Agronomy 11:2201

70. Lee K, Korani W, Bentz PC, Pokhrel S, Ozias-Akins P, Harkess A, Vaughn J, Clevenger J (2025) Long-Read Low-Pass Sequencing for High-Resolution Trait Mapping. bioRxiv:2025.2001.2009.632261

71. Meadows JRS, Kidd JM, Wang G-D, Parker HG, Schall PZ, Bianchi M, Christmas MJ, Bougiouri K, Buckley RM, Hitte C, Nguyen AK, Wang C, Jagannathan V, Niskanen JE, Frantz LAF, Arumilli M, Hundi S, Lindblad-Toh K, Ginja C, Agustina KK, André C, Boyko AR, Davis BW, Drögemüller M, Feng X-Y, Gkagkavouzis K, Iliopoulos G, Harris AC, Hytönen MK, Kalthoff DC, Liu Y-H, Lymberakis P, Poulakakis N, Pires AE, Racimo F, Ramos-Almodovar F, Savolainen P, Venetsani S, Tammen I, Triantafyllidis A, vonHoldt B, Wayne RK, Larson G, Nicholas FW, Lohi H, Leeb T, Zhang Y-P, Ostrander EA (2023) Genome sequencing of 2000 canids by the Dog10K consortium advances the understanding of demography, genome function and architecture. Genome Biology 24:187

72. Zhang X, Zhang X, Wang L, Liu Q, Liang Y, Zhang J, Xue Y, Tian Y, Zhang H, Li N, Sheng C, Nie P, Feng S, Liao B, Bai D (2023) Fine mapping of a QTL and identification of candidate genes associated with cold tolerance during germination in peanut (*Arachis hypogaea* L.) on chromosome B09 using whole genome re-sequencing. Frontiers in Plant Science 14:1153293

